## Supplementary material for "Multiomics mapping and characterization of cellular senescence in aging human skeletal muscle uncovers a novel senotherapeutic for sarcopenia": Suppl. Figure S1-S8

**Suppl. Fig. S1 Li Y & Li CH et al.**

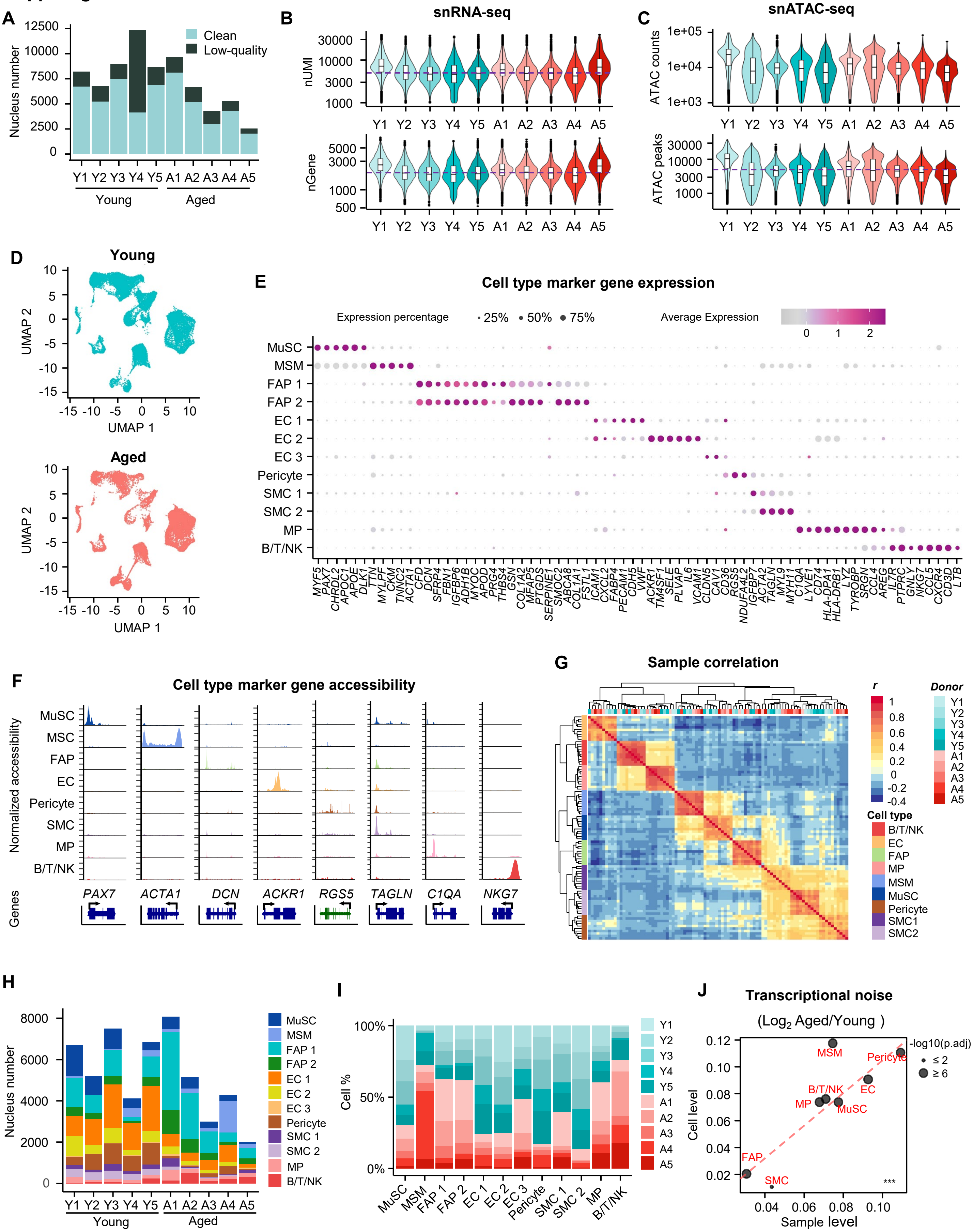

**Suppl. Fig. S2 Li Y & Li CH et al.**

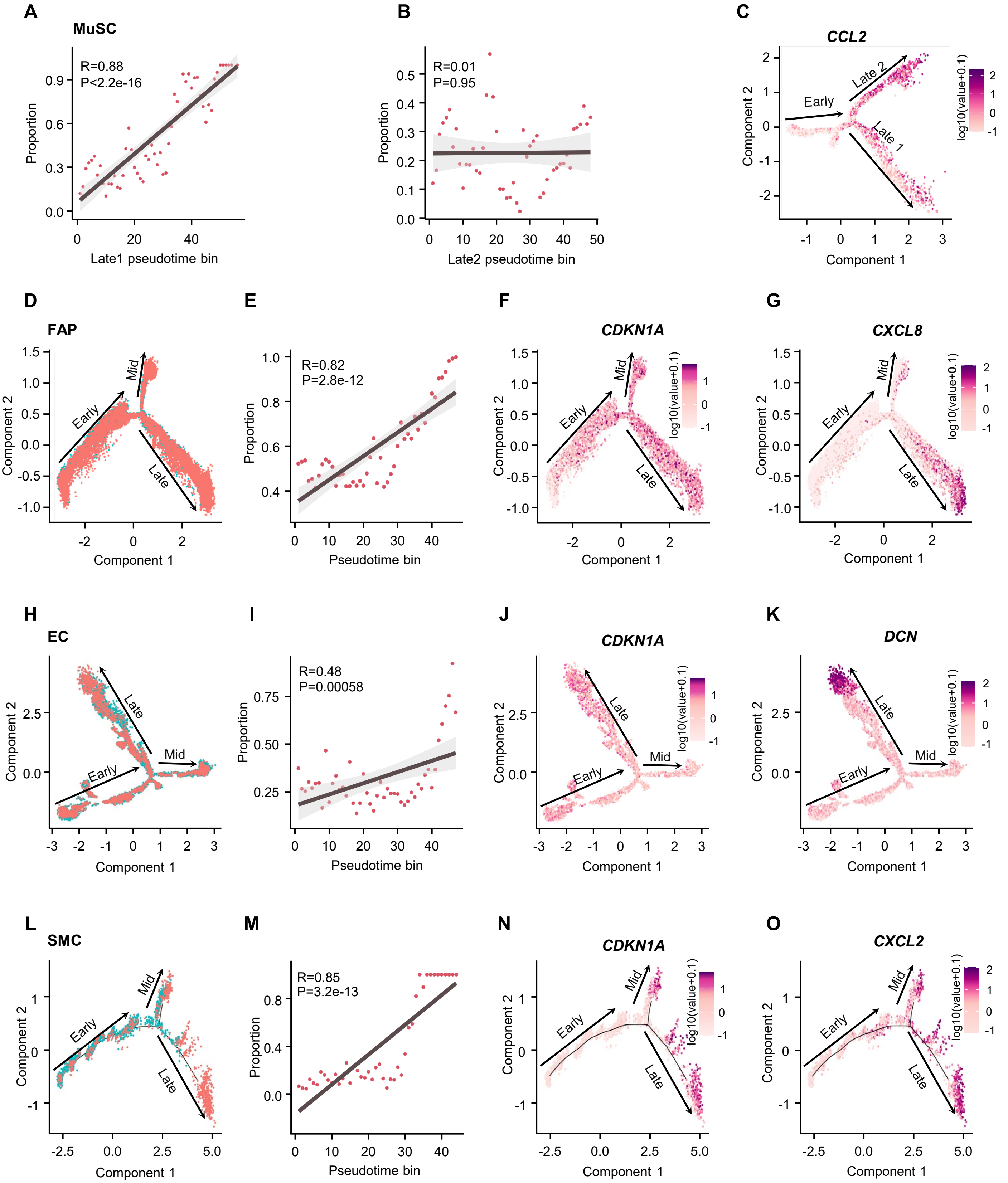

Suppl. Fig. S3 Li Y & Li CH et al.

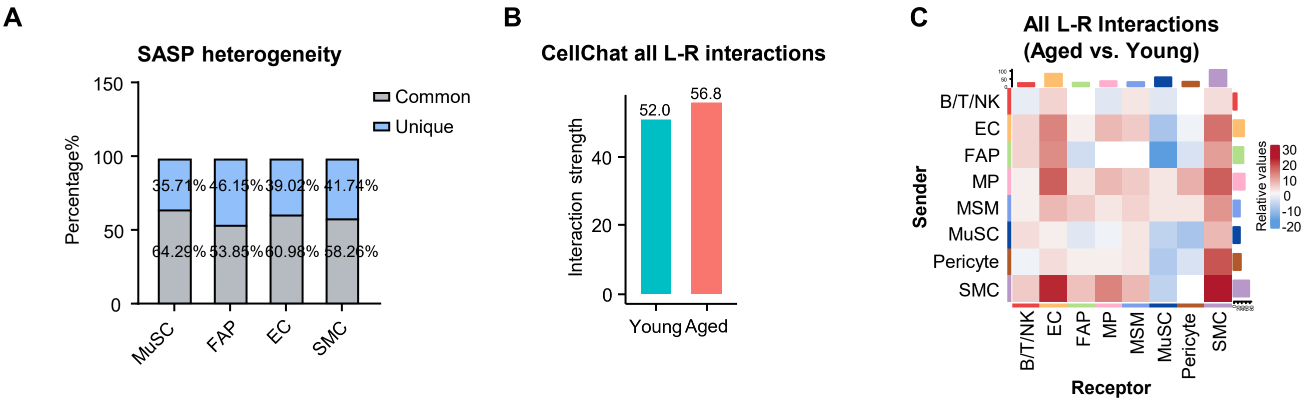

**Suppl. Fig. S4 Li Y & Li CH et al.**

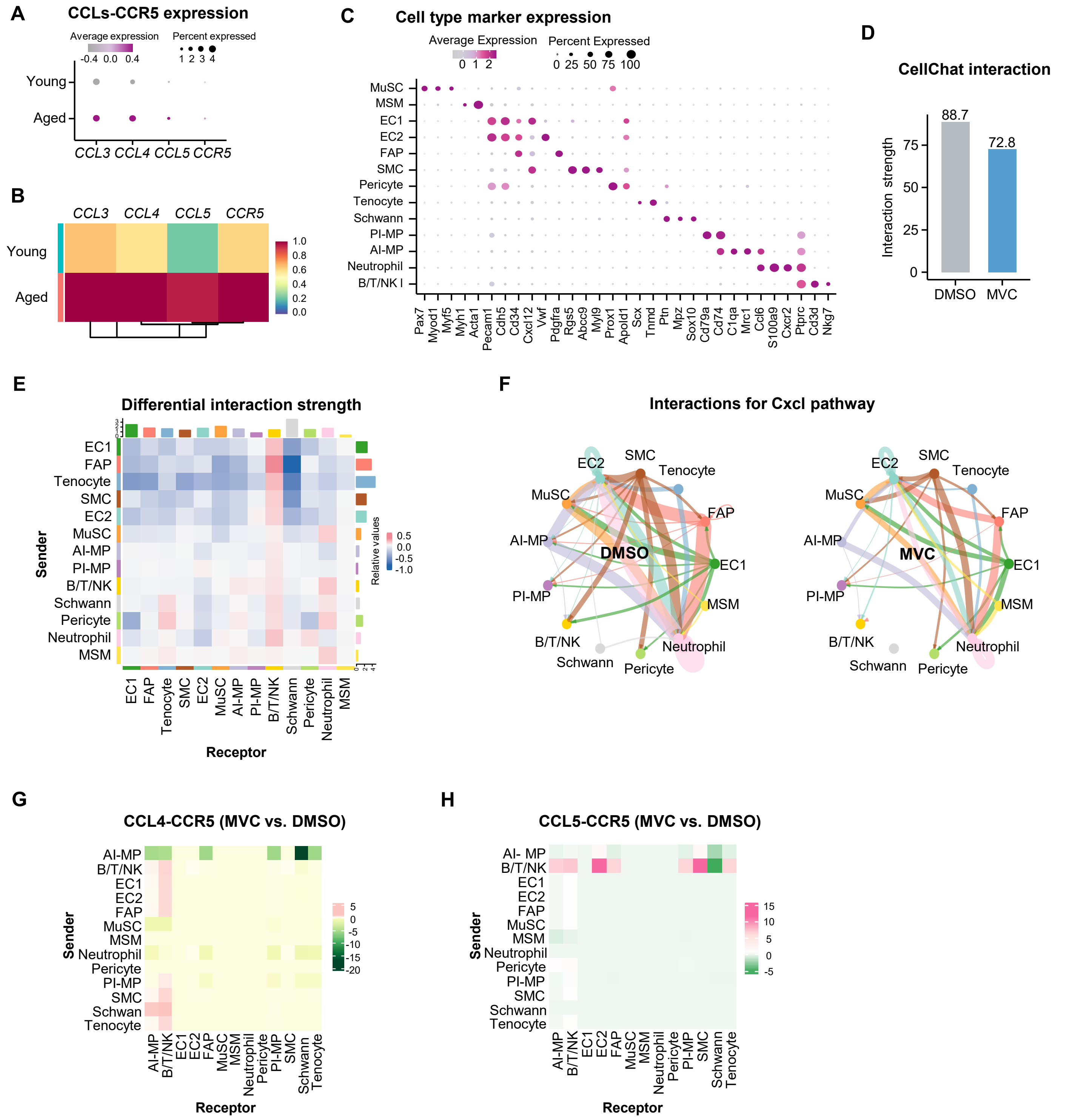

**Suppl. Fig. S5 Li Y & Li CH et al.**

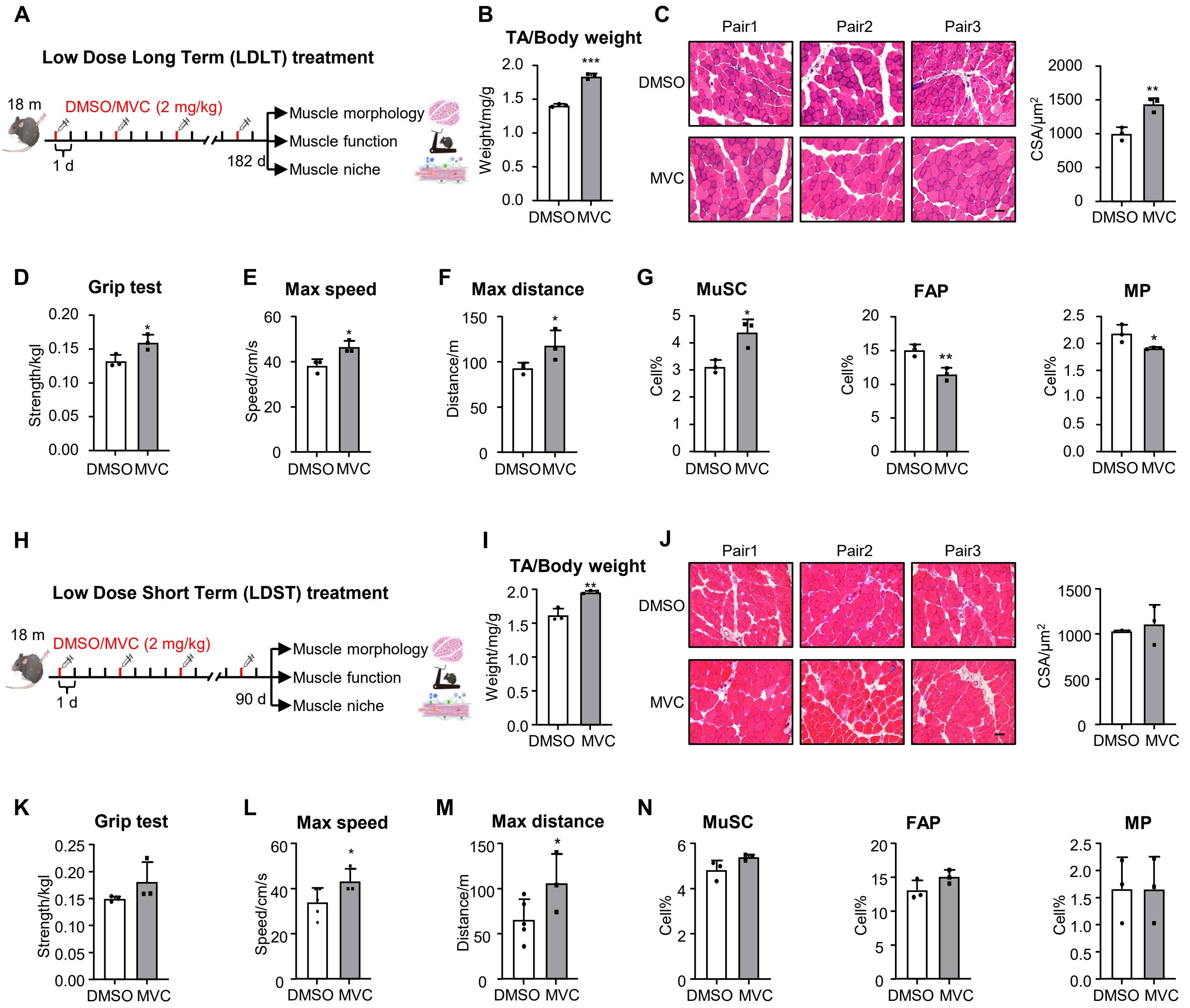

Suppl. Fig. S6 Li Y & Li CH et al.

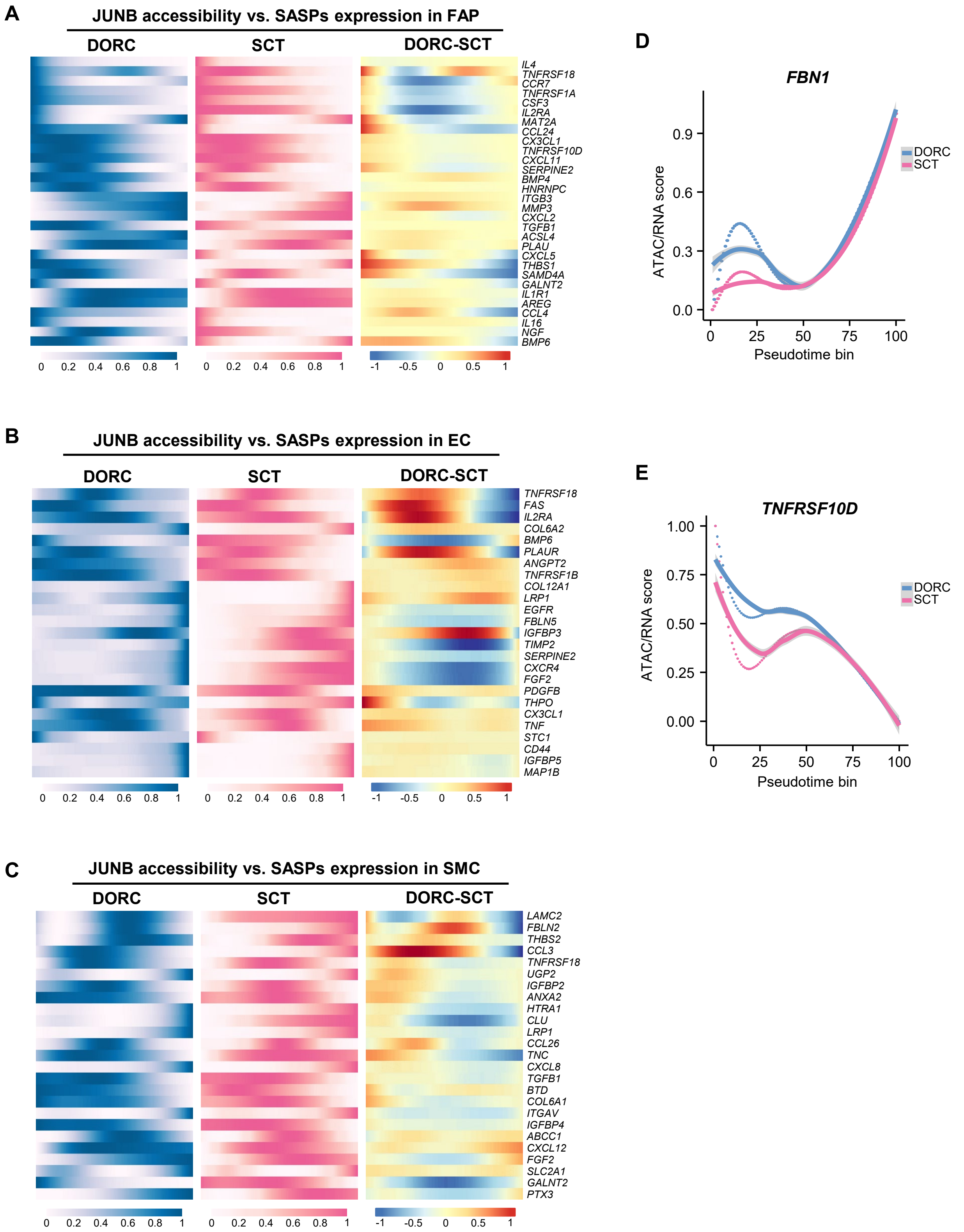

Suppl. Fig. S7 Li Y & Li CH et al.

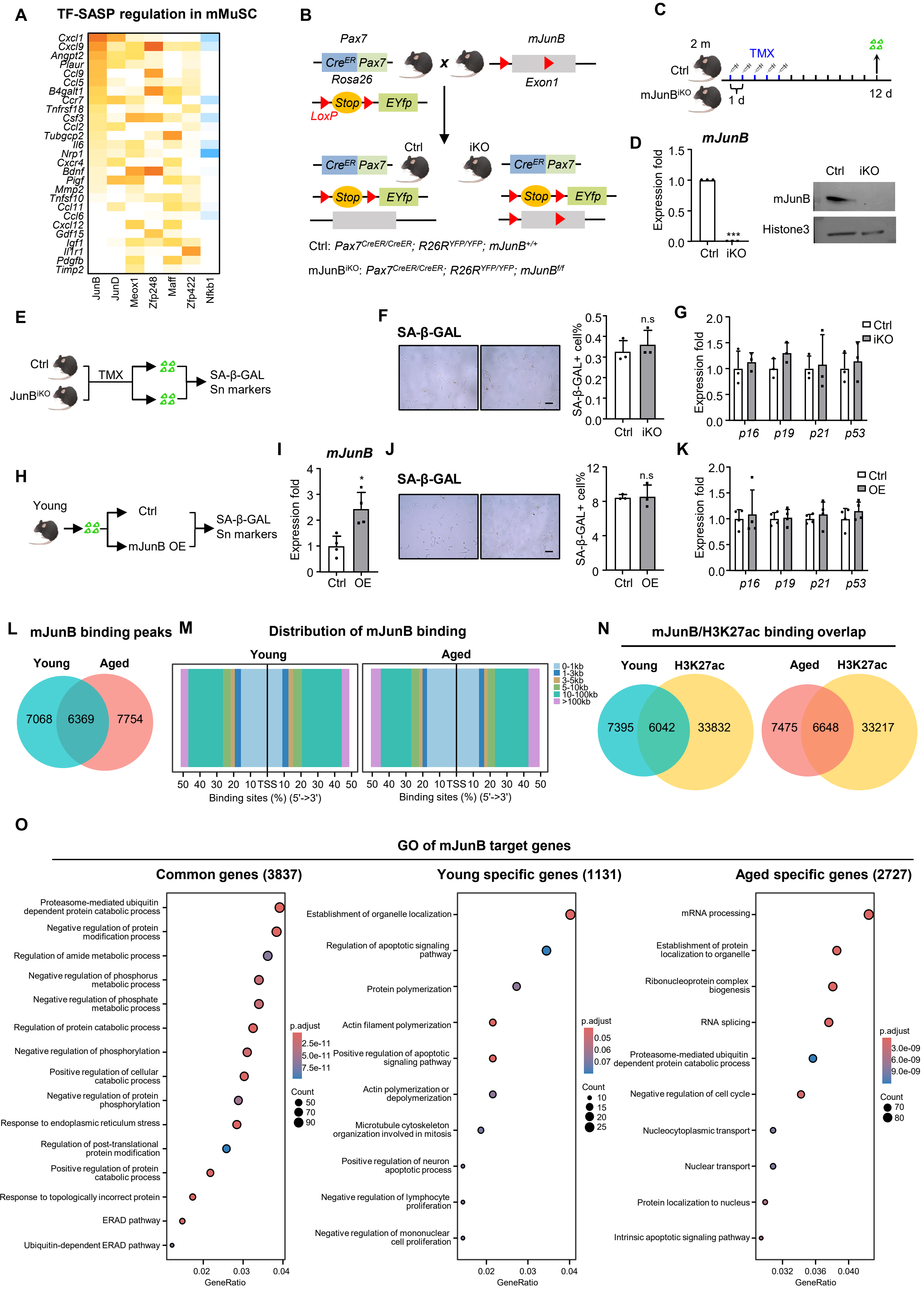

**Suppl. Fig. S8 Li Y & Li CH et al.**

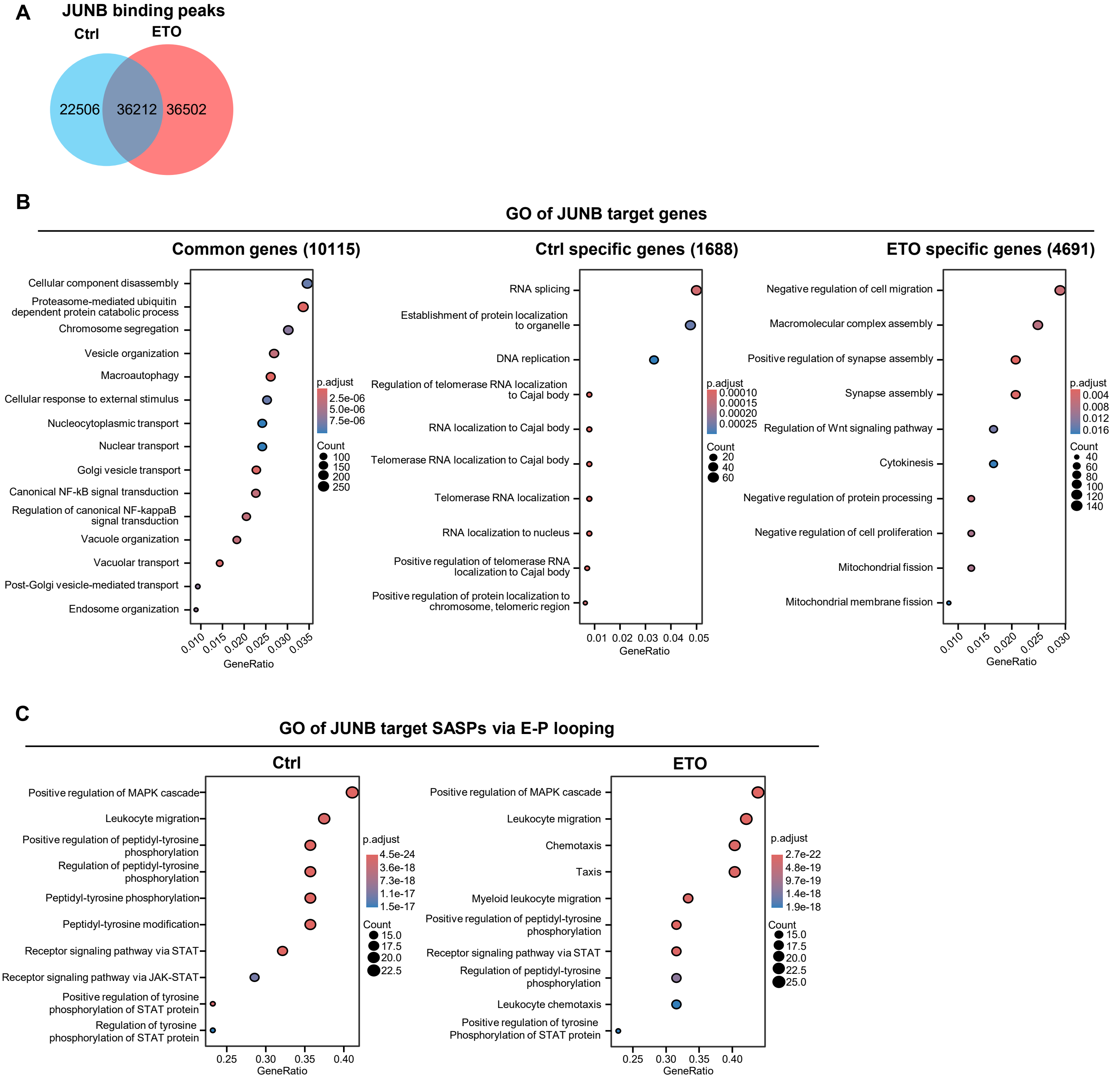
