## Supplementary material for "Multiomics mapping and characterization of cellular senescence in aging human skeletal muscle uncovers a novel senotherapeutic for sarcopenia": Suppl. information

**Inventory of Supplementary Information**

### 1. Supplementary Figures

Suppl. Fig. S1. Multimomics mapping of senescence atlas in aging human muscle.

Suppl. Fig. S2. Heterogeneity and dynamics of cellular senescence in aging human muscle.

Suppl. Fig. S3. SASP profiling and function in senescent cells.

Suppl. Fig. S4. Maraviroc is a potential senotherapeutic for sarcopenia.

Suppl. Fig. S5. Additional treatment regime of Maraviroc for sarcopenia.

Suppl. Fig. S6. Defining TFs governing senescence state and SASP induction in human muscle.

Suppl. Fig. S7. JUNB activates SASP induction in senescent MuSCs via enhancer regulation.

Suppl. Fig. S8. JUNB inhibition rejuvenates senescent human fibroblasts.

### 2. Supplementary Tables

Suppl. Table S1. Human skeletal muscle donor information

Suppl. Table S2. snRNA-seq analysis of the single-nuclei multiome data in aging human skeletal muscle

Suppl. Table S3. DEGs and GO analysis from snRNA-seq data

Suppl. Table S4. Analysis of SASP atlas and cell-cell communications in aging human skeletal muscle

Suppl. Table S5. Single-cell RNA-seq profiling in DMSO and MVC treatment muscle

Suppl. Table S6. Bulk RNA-seq analysis of MuSCs after MVC treatment

Suppl. Table S7. snATAC- and snRNA-seq analysis of TF-gene regulation

Suppl. Table S8. JunB CUT&RUN-seq analysis in mouse MuSCs

Suppl. Table S9. JUNB CUT&RUN-seq analysis in human IMR-90 cells

Suppl. Table S10. Sequences of oligonucleotides used in the study

### 3. Supplementary figure legends

**Suppl. Fig. S1. Multiomics mapping of senescence atlas in aging human muscle. A** Bar plot showing nucleus number from each human young (Y) or aged(A) donor skeletal muscle sample. **B** Box plot showing the unique molecular identifiers (UMIs) (top) and the number of genes (bottom) detected in each nucleus by analyzing snRNA-seq data. The dashed lines indicate 5000 UMIs and 2000 genes, respectively. **C** Box plot showing the number of ATAC counts and the number of peaks detected in each nucleus by analyzing the snATAC-seq data. The dashed line indicates the level of 5000 ATAC peaks. **D** UMAP visualization of young and aged groups showing no obvious batch effect. **E** Dot plot showing the snRNA-seq derived normalized expression levels of representative marker genes for each cell (sub)type. **F** snATAC-seq derived chromatin accessibility of representative marker genes (indicated below) for each cell type. **G** Heatmap showing the Pearson correlation of pseudo-bulked cell type-specific expression profiles among samples. **H** Bar plot showing the number of nuclei of each cell (sub)type captured in each of the young or aged samples. **I** Stacked bar plot showing the percentage of each sample in each of the 12 cell (sub)types. Colors used for indicating cell type are consistent with main figures. **J** Scatter plot showing the $\log_2$ ratio of transcriptional noise between aged and young samples as calculated using sample averages ($n$ = 10) and single cells on the X and Y axes, respectively. The dot size correlates with the negative $\log_{10}$ adjusted p-value of the cell type-level differential transcriptional noise analyzed using Wilcoxon test and the red dashed line corresponds to the robust F-test for comparing two linear models.

**Suppl. Fig. S2. Heterogeneity and dynamics of cellular senescence in aging human muscle. A** Scatter plot showing the proportion of aged MuSCs along the Early-Late1 pseudotime in 52 time bins (sized 0.1 per time bin). Pearson correlation of the proportion of aged nuclei and pseudotime: $R = 0.87$ and $P < 2.2 \times 10^{-16}$ (two-sided), with 95% CI (gray) shown. **B** Scatter plot showing the proportion of aged MuSCs along the Early-Late2 pseudotime in 47 time bins (sized 0.1 per time bin). Pearson correlation: $R = 0.01$ and $P = 0.95$ (two-sided), with 95% CI (gray) shown. **C** Expression level of

*CCL2* gene projected into discriminative dimensionality reduction (DDR) tree visualization of MuSCs. **D** DDR tree visualization of FAP trajectory with mapping of age group information. **E** Scatter plot showing the proportion of aged FAPs along the Early-Late pseudotime bins (47 time bins, sized 0.15 per time bin). Pearson correlation: $R = 0.82$ and $P = 2.8 \times 10^{-12}$ (two-sided), with 95% CI (gray) shown. **F-G** Expression levels of *CDKN1A* and *CXCL8* genes projected into DDR tree visualization of FAPs. **H** DDR tree visualization of EC trajectory with mapping of age group information. **I** Scatter plot showing the proportion of aged ECs along the Early-Late pseudotime bins (47 time bins, sized 0.20 per time bin). Pearson correlation: $R = 0.48$ and $P = 5.8 \times 10^{-4}$ (two-sided), with 95% CI (gray) shown. **J-K** Expression levels of *CDKN1A* and *DCN* gene projected into DDR tree visualization of ECs. **L** DDR tree visualization of SMC trajectory with mapping of age group information. **M** Scatter plot showing the proportion of aged SMCs along the Early-Late pseudotime bins (44 time bins, sized 0.20 per time bin). Pearson correlation: $R = 0.85$ and $P = 3.2 \times 10^{-13}$ (two-sided), with 95% CI (gray) shown. **N-O** Expression levels of *CDKN1A* and *CXCL2* genes projected into DDR tree visualization of SMCs.

**Suppl. Fig. S3. SASP profiling and function in senescent cells.** **A** Bar plot showing the ratios of common and cell type specific SASPs in MuSCs, FAPs, ECs and SMCs. **B** Bar plot comparing the interaction strength of all Ligand (L)-Receptor (R) mediated intercellular communications in aged vs. young group. **C** Heatmap showing differential number of all L-R mediated interactions between two cell types. Red/Blue represents increased/decreased signaling in the aged vs. young. The colored bar plot on the top or right represents the incoming/outgoing signaling setting each cell type as receptor/sender (sum of column/row of displayed values).

**Suppl. Fig. S4. Maraviroc is a potential senotherapeutic for sarcopenia.** **A** Dot plot showing snRNA-seq derived expression of *CCL3*, *CCL4*, *CCL5* and *CCR5* (*CCR5* axis genes) in all human muscle mononuclear cells. **B** Heatmap showing the pseudo-bulked expression level of *CCR5* axis genes in all mononuclear cells of young or aged muscle. **C** Dot plot showing the expression signatures of representative marker genes for each cell type. **D** Bar plot showing the interaction strength of intercellular communications calculated by CellChat in DMSO or MVC group. **E** Heatmap showing differential number of SASP-mediated interactions between two cell types. Red/Blue represents increased/decreased signaling in the MVC vs. DMSO. The colored bar plot on the top or right represents the incoming/outgoing signaling setting each cell type as receptor/sender (sum of column/row of displayed values). **F** Circle plot showing the signal strength change by aggregating all L-R pairs within Cxcl pathway. The edge colors correspond to the sender cell types, and the edge weights are proportional to the interaction strength. **G-H** Heatmap showing the interaction frequency of Ccl4-Ccr5 and Ccl5-Ccr5 across different cell types in MVC vs. DMSO.

**Suppl. Fig. S5. Additional treatment regime of Maraviroc for sarcopenia.** **A** Schematic of low dose long term (LDLT) treatment/assessment regime of MVC effect in aging muscle. **B** The ratio of TA/body weight of the treated mice, n=3. **C** Left: H&E staining of tibialis anterior (TA) muscles collected from the above-treated mice. Right: Quantification of cross sectional areas (CSAs) of the H&E stained fibers, Scale bar: 50 µm, n=3. **D** The treated mice were subject to a grip strength meter for strength measurement, n=3. **E-F** The treated mice were subject to treadmill exercise and the maximal running speed and distance were recorded, n=3. **G** Flow cytometry detection of the percentages of MuSC, MP, and FAP populations in the treated mice, n=3. **H** Schematic of low dose short term (LDST) treatment/assessment regime of MVC effect in aging muscle. **I** The ratio of TA/body weight of the treated mice, n=3. **J** Left: H&E staining of tibialis anterior (TA) muscles collected from the above-treated mice. Right: Quantification of CSAs of the stained fibers, Scale bar: 50 µm, n=3. **K** The treated mice were subject to a grip strength meter for strength measurement, n=3. **L-M** The treated mice were subject to treadmill exercise; the maximal running speed and distance were recorded, n=3. **N** Flow cytometry detection of the percentages of MuSC, MP, and FAP populations in the treated mice, n=3. All the bar graphs are presented as mean + SD, Student's t-test was used to calculate the statistical significance (B-G, I-N): *p < 0.05, **p < 0.01, ***p < 0.001, n.s = no significance.

**Suppl. Fig. S6. Defining TFs governing senescence state and SASP induction in human muscle.** **A-C** Heatmaps highlighting smoothed normalized JUNB DORC accessibility, SCT-normalized RNA expression, and the level of difference (DORC-RNA) for JUNB-target SASP genes identified to be significantly associated with FAP (A), EC (B), and SMC (C) aging pseudotime. **D-E** JUNB Chromatin (DORC) versus normalized gene expression (RNA) dynamics of SASP genes *FBN1* (D) and *TNFRSF10D* (E) with respect to MuSC aging pseudotime. The dotted line represents a LOESS fit to the JUNB DORC accessibility/gene expression dynamics along pseudotime in a gene-wise manner, derived from smooth spline curves fitted to the values obtained from 100 pseudotime bins.

**Suppl. Fig. S7. JUNB activates SASP induction in senescent MuSCs via enhancer regulation. A** Heatmap showing snATAC-seq detected DORC regulation scores for top-ranked TF-SASP association in mouse MuSCs. **B** Breeding scheme for generating inducible JunB conditional knock out (JunB$^{iKO}$: $Pax7^{CreERT2/R26YFP}$; $JunB^{f/f}$) and Control (Ctrl: $Pax7^{CreERT2/R26Yfp}$; $JunB^{+/+}$) mice. **C** Schematic of the experimental design for inducing JunB specific knocking out in MuSCs by Tamoxifen (TMX) injection. **D** RT-qPCR and western blot detection of the RNA and protein expression levels of JunB in Ctrl vs. iKO MuSCs, n=3. **E** Schematic of the experimental design for analyzing senescent MuSCs from Ctrl and JunB-iKO mice. **F** SA-β-GAL staining showing the percentage of senescent MuSCs in Ctrl vs. iKO. Scale bar: 50 μm, n=3. **G** RT-qPCR detection of the expression levels of senescent marker genes in Ctrl and iKO MuSCs, n=3. **H** Schematic of the experimental design for analyzing senescence in MuSCs with JunB overexpression. **I** RT-qPCR confirmation of the *JunB* overexpression after transfecting the JunB overexpressing or Ctrl plasmid into the MuSCs, n=3. **J** SA-β-GAL staining showing the senescent MuSCs in the above cells. Scale bar: 50 μm, n=3. **K** RT-qPCR detection of the expression levels of senescent marker genes in the above cells, n=3. **L** Pie chart showing the overlapping of JunB CUT&RUN-seq identified binding peaks in young and aged MuSCs. **M** Stacked bar chart showing the feature distribution of JunB binding sites based on the distance to TSS sites. **N** Pie chart showing the overlapping of JunB binding with H3K27ac in both young and aged MuSCs. **O** GO analysis of JunB target genes in young and aged MuSCs. The common and young or aged unique genes were shown respectively. All the bar graphs are presented as mean + SD, Student's t-test was used to calculate the statistical significance (D, F-G, I-K): *$p < 0.05$, **$p < 0.01$, ***$p < 0.001$, n.s = no significance.

**Suppl. Fig. S8. JUNB inhibition rejuvenates senescent human fibroblasts. A** Pie chart showing the overlapping of JUNB CUT&RUN-seq identified binding peaks in Ctrl and ETO-treated IMR-90 cells. **B** GO analysis of JUNB target genes in Ctrl and ETO-treated IMR-90. The common and unique genes are respectively shown. **C** GO analysis of JunB target SASPs regulated via E-P looping in Ctrl and ETO-treated IMR-90 cells.
